# Scheduled feeding improves sleep in a mouse model of Huntington’s disease

**DOI:** 10.1101/2024.05.04.592428

**Authors:** Emily Chiem, Kevin Zhao, Derek Dell’Angelica, Cristina A. Ghiani, Ketema N. Paul, Christopher S. Colwell

**Author notes:** Colwell and Paul are joint senior authors. Corresponding Author: Christopher S. Colwell Department of Psychiatry & Biobehavioral Sciences Semel Institute University of California Los Angeles Los Angeles, CA 90095 USA or Ketema N. Paul Department of Integrative Biology and Physiology Semel Institute University of California Los Angeles Los Angeles, CA 90095 USA.

## Abstract

Sleep disturbances are common features of neurodegenerative disorders including Huntington’s disease (HD). The sleep and circadian disruptions are recapitulated in animal models, and these models provide the opportunity to evaluate whether circadian interventions can be effective countermeasures for neurodegenerative disease. Time restricted feeding (TRF) interventions successfully improve activity rhythms, sleep behavior and motor performance in mouse models of HD. Seeking to determine if these benefits of scheduled feeding extend to physiological measures of sleep, electroencephalography (EEG) was used to measure sleep/wake states and polysomnographic patterns in adult mice (six mo-old) under TRF and *ad lib* feeding (ALF). With each diet, both male and female wild-type (WT) and bacterial artificial chromosome transgenic (BACHD) mice were evaluated. Our findings show that male, but not female, BACHD mice exhibited significant changes in the temporal patterning of wake and non-rapid eye movement (NREM) sleep. The TRF intervention reduced the inappropriate early morning activity by increasing NREM sleep in the male BACHD mice. In addition, the scheduled feeding reduced sleep fragmentation (# bouts) in the male BACHD mice. The phase of the rhythm in rapid-eye movement (REM) sleep was significantly altered by the scheduled feeding. The treatment did impact the power spectral curves during the day in male but not female mice. Sleep homeostasis, as measured by the response to six hours of gentle handling, was not altered by the diet. Thus, TRF improves the temporal patterning and fragmentation of NREM sleep without impacting sleep homeostasis. This work adds critical support to the view that sleep is a modifiable risk factor in neurodegenerative diseases.

---

Sleep disturbance is a common feature of neurodegenerative diseases, such as Huntington’s disease (HD) (Colwell, 2021; Voysey et al., 2021). Huntington’s disease is caused by an abnormal CAG repeat expansion within the huntingtin (Htt) gene, which leads to widespread physiological disruption (Tabrizi et al., 2020). Sleep questionnaires find that HD patients commonly experience insomnia, daytime sleepiness, delayed sleep onset, and frequent nighttime awakenings (Goodman et al., 2011; Herzog-Krzywoszanska and Krzywoszanski, 2019; Tanigaki et al., 2020; Ogilvie et al., 2021). Smaller scale polysomnography (PSG) studies have also uncovered delayed sleep onset, increased sleep fragmentation as well as decreased slow-wave sleep (Arnulf et al., 2008; Cuturic et al., 2009; Goodman et al., 2011; Lazar et al., 2015). Broadly, these sleep/wake cycle disturbances described in the HD carriers are recapitulated in mouse models. The R6/2 mouse model of HD exhibits increased activity during the day and sleep fragmentation (Morton et al., 2005; Kantor et al., 2013) and the Q175 model has been shown to display increased wakefulness and decreased NREM sleep during the light phase (Loh et al., 2013; Fisher et al., 2016). In addition, the BACHD mouse model of HD exhibits sleep/wake architecture disruptions (Kudo et al., 2011), with sex differences in these disturbances (Chiem et al., 2024). Overall, there is comprehensive evidence of abnormal electroencephalography (EEG)-defined sleep architecture in mouse (Kantor et al., 2013; Fisher et al., 2013; Lebreton et al., 2015, Fisher et al., 2016) and sheep (Schneider et al., 2021, Vas et al., 2021) models of HD. These studies consistently found an early and progressive deterioration of both sleep architecture and behavior.

Diurnal rhythms in the sleep/wake cycle are generated, in part, by the circadian timing system with a circadian clock in the suprachiasmatic nucleus (SCN). In HD patients, the SCN shows signs of extensive degeneration (van Wamelen et al., 2014). In mouse models, there are reductions in the expression of the neuropeptide vasoactive intestinal polypeptide (VIP) and its receptor VPAC2 within the SCN (Fahrenkrug et al. 2007; Kuljis et al., 2016) as well as reductions in the neural activity rhythms that are the hallmark of SCN function (Kudo et al., 2011; Kuljis et al., 2016; Kuljis et al., 2018). Therefore, in the case of HD and perhaps other neurodegenerative models, the ideal intervention would be effective even with a compromised SCN. The daily feed/fast cycle is a powerful regulator of the circadian system that functions even when the SCN is damaged (Stephan, 1983; 1989; Angeles-Castellanos et al., 2010; Mistlberger, 2011). In early work with the R6/2 model, food entrainment aided in the maintenance of body temperature rhythms and improved locomotor behavior (Maywood et al., 2010; Skillings et al., 2014). A time-restricted feeding (TRF) protocol has been shown to improve locomotor activity, sleep behavioral patterns, and heart rate variability in Q175 mice (Wang et al., 2018) and BACHD mice (Whittaker et al., 2018). This prior work has not examined the possible impact of TRF on EEG-defined sleep. Thus, in the present study, we utilized the BACHD mouse model of HD to examine the impact of TRF on sleep/wake architecture, EEG spectral power, and the homeostatic response to sleep deprivation.

## Methods

### Animals

All the experimental protocols used to collect the data for the present report were approved by the UCLA Animal Research Committee and followed the guidelines and recommendations for animal use and welfare set by the UCLA Division of Laboratory Animal Medicine and National Institutes of Health. The BACHD mouse model of HD contains a human mutant Htt gene encoding 97 glutamine repeats (Gray et al., 2008). BACHD females backcrossed on a C57BL/6J background were bred in-house with C57BL/6J (WT) males from the Jackson Laboratory in order to obtain male and female offspring, either WT or heterozygous for the BACHD transgene. The WT littermates were used as WT controls in this study. There were four groups of mice used in this study including both male (WT, n=21; BACHD, n=15) and female (WT, n=13; BACHD, n=16) mice in order to further our knowledge on the sex difference. Mice with severe artifacts in their recordings were excluded from analysis including three WT and one BACHD. Animals were group housed (4 per cage), and entrained to a 12:12 LD cycle, in sound-proof, humidity-controlled chambers until experimentation began.

### Surgery

All animals were surgically implanted at 12 weeks with EEG and electromyograph (EMG) electrodes for polysomnography recordings. A prefabricated headmount (Pinnacle Technologies, Lawrence, KS) was used to position three stainless-steel epidural screw electrodes. The first electrode (frontal-located over the frontal cortex) was placed 1.5 mm anterior to bregma and 1.5 mm lateral to the central suture. The second two electrodes (interparietal-located over the visual cortex and common reference) were placed 2.5 mm posterior to bregma and 1.5 mm on either side of the central suture. The resulting two leads (frontal-interparietal and interparietal-interparietal) were referenced contralaterally. A fourth screw served as ground. Silver epoxy was used to aid electrical continuity between the screw electrode and headmount. Stainless-steel teflon-coated wires were inserted bilaterally into the nuchal muscle to record EMG activity. The headmount was secured to the skull with dental acrylic.

### EEG/EMG recording

Mice were placed in sound-proof sleep-recording chambers and connected to a lightweight tether attached to a low-resistance commutator mounted over the cage (Pinnacle Technologies). The animals were allowed free range of movement throughout the cage while being tethered and given one week to acclimate to the tether and recording chambers. EEG and EMG recordings began at zeitgeber time (ZT) 0 (light onset) and continued for 24h. Data acquisition was performed on a personal computer running Sirenia Acquisition software (Pinnacle Technologies), a software system specific to rodent polysomnographic recordings.

EEG signals were low-pass filtered with a 40-Hz cutoff and collected continuously at a sampling rate of 400 Hz. After data collection, waveforms were scored by the same trained operator as wake (low-voltage, high-frequency EEG; high-amplitude EMG), NREM sleep (high voltage, mixed frequency EEG; low-amplitude EMG), or REM sleep (low-voltage EEG with a predominance of theta activity (5-7Hz); low amplitude EMG). EEG epochs containing artifact due to scratching, moving, eating, or drinking were excluded from analysis. Recordings were scored in 10-sec epochs. The operator was masked to the sex and genotype of the mice.

### Signal Analysis

Spectral analysis was performed on the frontal-interparietal lead. Power spectral analysis was performed by applying a fast Fourier transform (FFT) to raw EEG waveforms. Absolute power (μV^2^) was analyzed in 0.1 Hz bins across the entire power spectrum (0-40 Hz) using Sirenia Sleep Pro software (Pinnacle Technologies). Relative power in the delta (0.5-4 Hz), theta (5-7 Hz), alpha (8-12 Hz), beta (14-20 Hz), and gamma (20-40 Hz) bands was measured across the 24-hr period. Relative NREM delta power during recovery (ZT6-24) was normalized to the average 24-hour baseline NREM delta power for each animal. Sleep fragmentation was measured by the number of NREM sleep bouts and duration of N sleep bouts (sec).

### Total sleep deprivation

Immediately following a 24-hr baseline recording, mice underwent 6-hrs of total sleep deprivation (SD) using a gentle-handling protocol, which includes cage tapping, introduction of novel objects, and delicate touching when mice displayed signs of sleep onset. SD began at the onset of the light phase in a 12hr:12hr LD cycle. Recordings continued for an 18-hr of recovery opportunity following the period of forced wakefulness.

### Time restricted feeding (TRF)

Male and female WT and BACHD mice (3-4 months old) were exposed to one of two feeding conditions for 1 month: *ad libitum* feeding (ALF) or feeding restricted to 6 hrs during the middle of the active phase (ZT15-21). All mice had *ad lib* access to water. Mice were singly housed, and the bedding was changed twice a week as mice are coprophagic. For the first three weeks of TRF, mice were housed in cages with a custom-made programmable food hopper that controls food access. At the start of the fourth week of TRF, mice were connected to tethers and allowed one week to acclimate prior to EEG recordings. TRF was performed manually during this time, as the programmable food hoppers did not fit in the recording cages. Control mice were singly housed and given *ad lib* access to food and water.

### Statistical Analysis

For the analysis of the data generated in this study, we used a three-way ANOVA to analyze the effects of three independent variables (factors: genotype, treatment and sex or time) on dependent variables measured by the EEG. We evaluated the main effects of each factor as well as any interaction between the factors. When appropriate, multiple comparison post-hoc tests were used to assess significant differences. In addition, we also used a linear mixed effects (LME) model to examine the spectral data. Each of the two methods serve different purposes and has different assumptions. The LME offers particular advantages for repeated measures data while better accounting for subject-specific variability and the correlated structure of the data. The LME model incorporates fixed effects (genotype, treatment, sex, and time) as well as helping us understand the extent to which variability in the outcome is explained by the fixed and random effects. For the spectral curve analysis, we included spectral power values (Hz) as the dependent variable and fixed effects of genotype, time of day, treatment, sex, and frequency. Each animal was included as a random effect. The model specifications were as follows: power values ∼ Genotype * Time * Treatment * Sex * Frequency + (1 | Animal ID).

The LME analyses was conducted using the *lme4* package in R version 4.2.1 (Bates et al., 2015) while the ANOVAs were conducted using SigmaPlot (version 14.5; SYSTAT Software, San Jose, CA). All values are reported as group mean ± standard error of the mean (SEM). Each animal was considered an experimental unit. Since we expected a normal distribution, we defined an outlier as a value that falls 1.5 standard deviations above or below the mean of the values. As mentioned above, a few mice were removed from the data set for technical reasons as the artifacts prevented the determination of the clear signal. The waveforms under baseline or after SD were analyzed by three-way analysis of variance (ANOVA) with genotype, sex, and time as factors. The total time spent in each state (wake, REM and NREM sleep), the amplitude or phase of the rhythms, and response to SD were analyzed by two-way ANOVA with genotype (WT, BACHD) and sex (male, female) as factors. The Holm-Sidak multiple comparison test was used when appropriate. Between-group differences were determined significant if p < 0.05.

To evaluate the rhythmicity of individual animals, their wake/REM/NREM measures for 24-hrs were analyzed using the Acro program (https://www.circadian.org/softwar.html) by Dr. R. Refinetti to determine goodness of fit (0-1.0) with 1 being the worst. This program measures the similarity between the cosine of the angle between the vectors of the empirical data and a fitted cosine wave.

## Results

### Wake rhythms

In this study, we sought to use EEG measurements to test the hypothesis that TRF would impact daily rhythms of sleep/wake architecture and whether there were sex differences in the response. First, we examined the rhythm in wake looking at 2-hr bins across the 24 hrs cycle under ALF and TRF (**Fig. 1A, B**). We analyzed the waveforms using a three-way ANOVA with genotype, treatment, and time as factors for male and female mice. Time was highly significant for both sexes while the male mice also exhibited significant differences by genotype (**Table 1**). For both sexes, there was a significant interaction between treatment and time (**Table 1**). The elevated early morning wakefulness characteristic of the male BACHD was corrected by the TRF intervention (**Fig. 1A**). There were no significant differences in the peak phase of the wake rhythms between groups (**Fig. 1C, D**; **Table 2**). Still, the male, but not the female, BACHD exhibited a great deal of variability in phase (**Table 3**) compared to the WT, which was corrected by TRF. As measured by a cosine analysis, the percentage of mice with a significant diurnal rhythm in wake was increased by the treatment (**Fig. 1E, F**; **Table 4**). We quantified the amplitude of day/night difference as a ratio of minutes of wake in the night over the minutes measured during the day. TRF increased the amplitude of the rhythms with the strongest effects in the male BACHD (**Fig. 1G**; **Table 2**). The male BACHD mice exhibited more wake during the day (resting period) and less during the night (active period) and this difference corrected by TRF (**Fig. 1H**). The total amount of wake did not vary with genotype, sex, or feeding schedule (**Fig. 1I**, **Table 2**). Thus, as measured by EEG, TRF regularized the rhythm in the wake state in the male BACHD.

**Fig 1.**
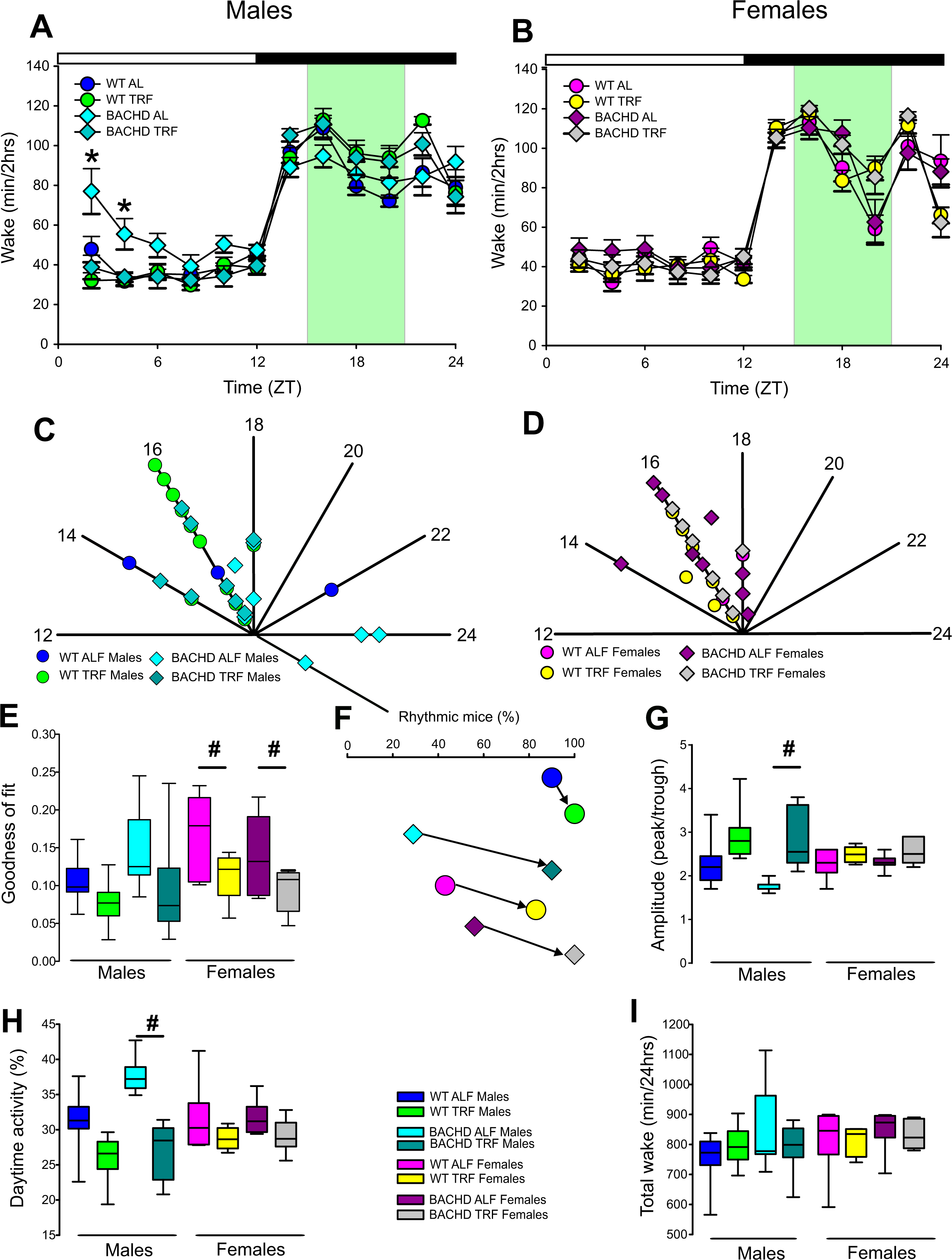
Temporal Pattern of Wake in BACHD and WT mice under TRF and ALF. EEG recordings were conducted over 24-hr in undisturbed mice held in a 12:12 light/dark (LD) cycle. Waveforms showing the daily rhythms of wake in male (**A**) and female (**B**) WT and BACHD mice plotted in 2-hrs bins. The BACHD males displayed increased wake during the lights-on (inactive) phase. This increase was corrected by the TRF intervention. Data are presented as mean ± SEM. Time points in which significant differences (P<0.05) were found between the groups using multiple comparison procedures (Holm-Sidak method) are indicated by asterisk. For this and the other figures, the timing of the light/dark cycle is shown by the bar at the top of the graph. The green-shaded area represents the time of schedules feeding for the TRF treated groups. The circles represent data from WT groups and the diamonds represent data from BACHD groups. Polar display of the peak phase of the rhythms of wake in male (**C**) and female (**D**) WT and BACHD mice. The numbers on the axis represent time (ZT) with ZT 12 indicating the time of lights-off. The male BACHD under ALF exhibited higher variability in peak phase than the other groups. (**E**) The vector of the diurnal rhythms in wake were fit to a cosine wave and the “goodness of fit” was determined for each animal. The lower the number the better the fit. (**F**) The percentage of mice exhibiting a significant diurnal rhythm for each group. The lines with arrow connect each genotype and sex under ALF and TRF. This display shows that the scheduled feeding increases the number of mice with rhythms in wake. The amplitude of the rhythms (**G**) as well as the % of wake during the day (**H**) and the total minutes of recorded wake (**I**) were displayed for each group. The TRF treatment increase the amplitude of the male BACHD rhythm as well as reduced the % activity in the day compared to BACHD mice under ALF. In this and other figures, the box plots visually show the distribution of numerical data and skewness by displaying the data quartiles and median.

**Table 1:** Analysis of waveforms by three-way ANOVA with genotype (WT, BACHD), treatment (ALF, TRF) and time (2 hr bins) as factors. Significant values are indicated in bold. Interactions between treatment and time are reported. Degrees of freedom are reported within parentheses, alpha=0.05. Bold type indicates statistical significance.

| State | Sex | Genotype | Time<br>(2 hr bins) | Treatment | Interaction<br>(treatment x Time) |
| --- | --- | --- | --- | --- | --- |
| Wake | males | **F<sub>(1,419)</sub> = 5.272, P < 0.022** | **F<sub>(11,419)</sub> = 109.768, P < 0.001** | F<sub>(1,419)</sub> = 0.270, P = 0.604 | **F<sub>(11,419)</sub> = 6.892, P < 0.001** |
|  | females | F<sub>(1,335)</sub> = 1.891, P = 0.170 | **F<sub>(11,335)</sub> = 117.352, P < 0.001** | F<sub>(1,335)</sub> = 0.180, P = 0.668 | **F<sub>(11,335)</sub> = 5.029, P < 0.001** |
| REM | males | F<sub>(1,419)</sub> = 1.141, P = 0.286 | **F<sub>(11,419)</sub> = 77.934, P < 0.001** | **F<sub>(1,419)</sub> = 5.559, P = 0.018** | F<sub>(11,419)</sub> = 1.526, P = 0.120 |
|  | females | **F<sub>(1,335)</sub> = 6.264, P = 0.013** | **F<sub>(11,335)</sub> = 125.641, P < 0.001** | F<sub>(1,335)</sub> = 0.126, P = 0.722 | **F<sub>(11,335)</sub> = 2.654, P = 0.003** |
| NREM | males | F<sub>(1,419)</sub> = 5.272, P = 0.022 | **F<sub>(11,419)</sub> = 109.768, P < 0.001** | F<sub>(1,419)</sub> = 0.270, P = 0.604 | **F<sub>(11,419)</sub> = 6.546, P = 0.011** |
|  | females | F<sub>(1,335)</sub> = 1.891, P = 0.170 | **F<sub>(11,335)</sub> = 117.352, P < 0.001** | F<sub>(1,335)</sub> = 0.184, P = 0.688 | **F<sub>(11,335)</sub> = 5.029, P < 0.001** |

**Table 2:** Analysis of key parameters of each state (wake, REM and NREM Sleep) by three-way ANOVA with genotype (WT, BACHD), sex (male, female)), and treatment (ALF, TRF) as factors. Degrees of freedom are reported within parentheses, alpha=0.05. Bold type indicates statistical significance.

| State |  | Genotype | Sex | Treatment |
| --- | --- | --- | --- | --- |
| Wake | Amplitude | F<sub>(1, 62)</sub> = 1.329, P = 0.254 | **F<sub>(1, 62)</sub> = 22.724, P < 0.001** | F<sub>(1, 62)</sub> = 0.096, P = 0.757 |
|  | Phase | F<sub>(1, 62)</sub> = 0.201, P = 0.656 | F<sub>(1, 62)</sub> = 0.955, P = 0.333 | F<sub>(1, 62)</sub> = 0.007, P = 0.932 |
|  | Day (%) | **F<sub>(1, 62)</sub> = 5.364, P = 0.024** | **F<sub>(1, 62)</sub> = 37.982, P < 0.001** | F<sub>(1, 62)</sub> = 0.105, P = 0.748 |
|  | Total | F<sub>(1, 62)</sub> = 2.656, P = 0.109 | F<sub>(1, 62)</sub> = 0.180, P = 0.673 | F<sub>(1, 62)</sub> = 2.112, P = 0.152 |
| REM | Amplitude | F<sub>(1, 56)</sub> = 0.089, P = 0.766 | F<sub>(1, 56)</sub> = 1.618, P = 0.209 | F<sub>(1, 56)</sub> = 0.444, P = 0.508 |
|  | Phase | F<sub>(1, 61)</sub> = 0.314, P = 0.577 | **F<sub>(1, 61)</sub> = 7.808, P = 0.007** | **F<sub>(1, 61)</sub> = 9.546, P = 0.003** |
|  | Night (%) | F<sub>(1, 61)</sub> = 1.160, P = 0.286 | F<sub>(1, 61)</sub> = 1.714, P = 0.196 | F<sub>(1, 61)</sub> = 0.154, P = 0.696 |
|  | Total | F<sub>(1, 61)</sub> = 3.730, P = 0.059 | F<sub>(1, 61)</sub> = 1.366, P = 0.248 | F<sub>(1, 61)</sub> = 0.988, P = 0.322 |
| NREM | Amplitude | F<sub>(1, 61)</sub> = 1.335, P = 0.253 | F<sub>(1, 61)</sub> = 0.991, P = 0.324 | F<sub>(1, 61)</sub> = 0.162, P = 0.689 |
|  | Phase | **F<sub>(1, 62)</sub> = 14.685, P < 0.001** | F<sub>(1, 62)</sub> = 3.662, P = 0.061 | **F<sub>(1, 62)</sub> = 6.440, P = 0.014** |
|  | Night (%) | F<sub>(1, 62)</sub> = 0.569, P = 0.812 | F<sub>(1, 62)</sub> = 2.706, P = 0.106 | F<sub>(1, 62)</sub> = 1.475, P = 0.230 |
|  | Total | F<sub>(1, 62)</sub> = 0.384, P = 0.538 | F<sub>(1, 62)</sub> = 3.554, P = 0.065 | **F<sub>(1, 62)</sub> = 5.970, P = 0.018** |

**Table 3:**
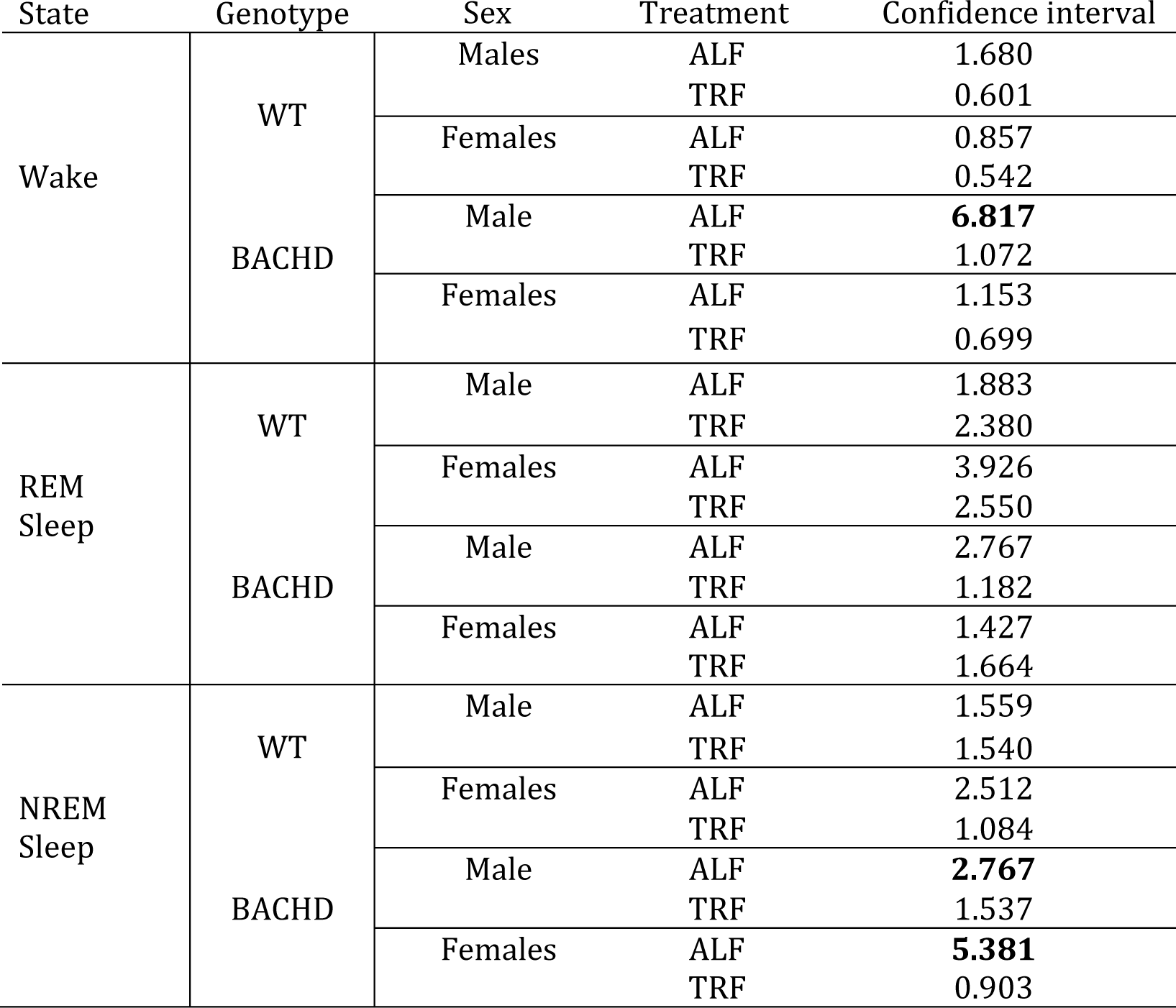
Confidence intervals of the peak phase of the EEG rhythms by genotype and sex. The BACHD males exhibited strikingly more variable measures for each state.

**Table 4:** Analysis of goodness of fit by three-way ANOVA with genotype (WT, BACHD), sex (males, females) and treatment (ALF, TRF) as factors. Significant values are indicated in bold. No significant interactions between the three factors were detected. Degrees of freedom are reported within parentheses, alpha=0.05. Bold type indicates statistical significance.

| State | Genotype | Sex | Treatment |
| --- | --- | --- | --- |
| Wake | $F_{(1,62)} = 0.382, P = 0.539$ | **$F_{(1,62)} = 4.246, P = 0.044$** | **$F_{(1,62)} = 20.117, P < 0.001$** |
| REM | $F_{(1,61)} = 0.032, P = 0.858$ | $F_{(1,61)} = 1.116, P = 0.298$ | **$F_{(1,62)} = 8.123, P = 0.006$** |
| NREM | $F_{(1,62)} = 0.049, P = 0.825$ | **$F_{(1,62)} = 9.428, P = 0.003$** | **$F_{(1,62)} = 20.611, P < 0.001$** |

### REM sleep rhythms

Next, we carried out a similar analysis on the temporal patterns of REM sleep. First, we examined the rhythm in REM sleep looking at 2-hr bins across the 24-hr cycle under ALF and TRF (**Fig. 2A, B**). We analyzed the waveforms using a three-way ANOVA with genotype, treatment, and time as factors for male and female mice. Time was highly significant for both sexes. The male mice also exhibited significant differences by treatment but not genotype while the female mice exhibited significant differences by genotype but not treatment (**Table 1**). For female mice, there was a significant interaction between treatment and time. The phase of REM sleep (**Fig. 2C, D**) was significantly affected by feeding schedules that differed by sex (**Table 2**). With REM sleep, we did not see a large increase in variability in the male BACHD by other measures, while the WT females exhibited the most variability in peak phase (**Table 3**). The percentage of mice with a significant diurnal rhythm in REM sleep was increased by the treatment in all but the WT males (**Fig. 2E, F**; **Table 4**). The amplitude of the rhythm in REM sleep or the total amount of REM sleep did not vary with genotype, treatment, or sex (**Fig. 2G, H, I**; **Table 2**). Thus, as measured by EEG, TRF altered the phase of the rhythm in REM sleep with the primary effect in females.

**Fig 2.**
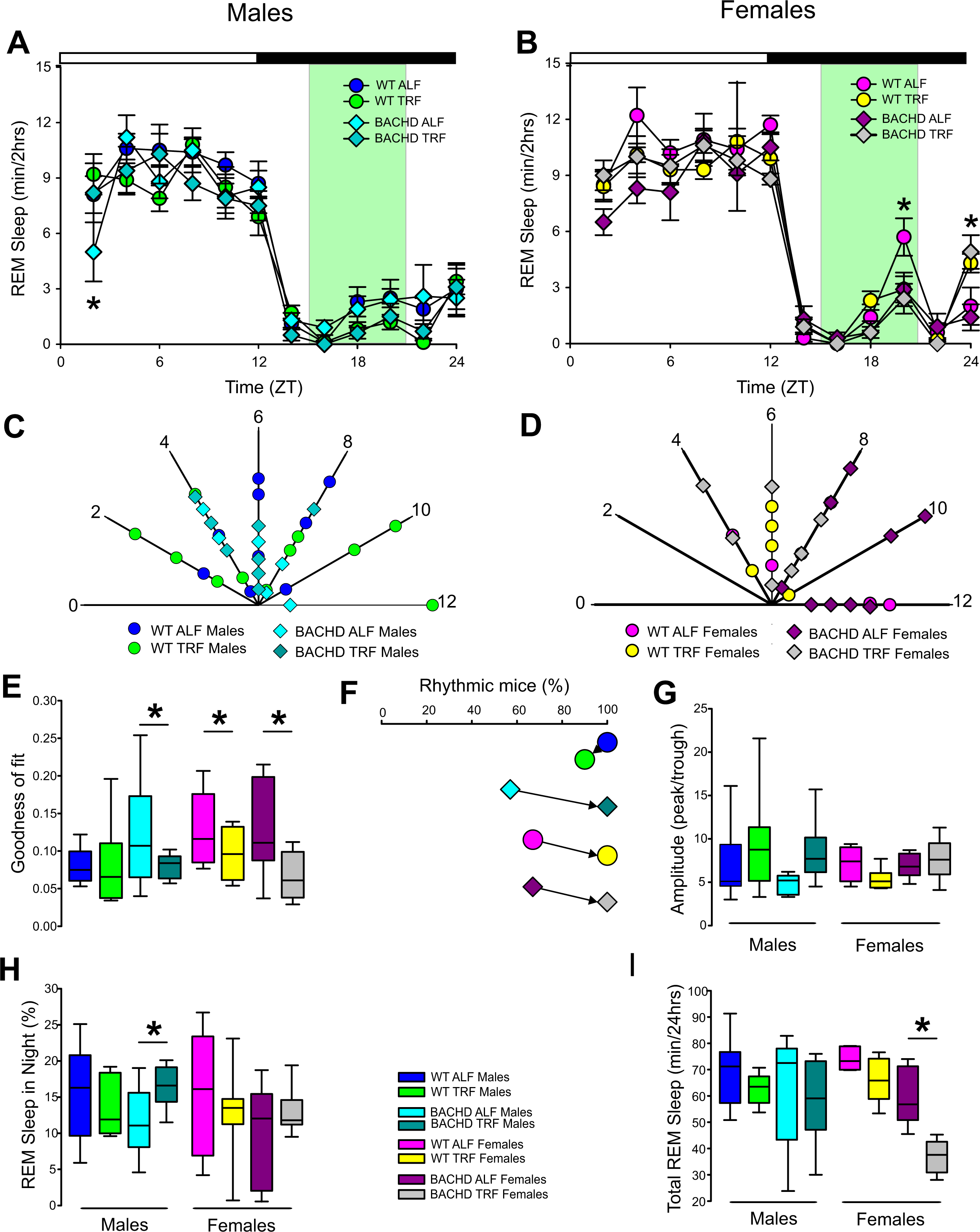
Temporal Pattern of REM sleep in BACHD and WT mice under TRF and ALF. EEG recordings were conducted over 24-hr in undisturbed mice held in a 12:12 LD cycle. Waveforms showing the daily rhythms of REM sleep in male (**A**) and female (**B**) WT and BACHD mice plotted in 2-hrs bins. The BACHD males displayed reduced REM sleep at ZT 2 and this reduction was corrected by the TRF intervention. In WT females, the intervention reduced REM sleep at ZT 20. Polar display of the peak phase of the rhythms of REM sleep in male (**C**) and female (**D**) WT and BACHD mice. There were not obvious differences between the groups in phase of REM sleep. The “goodness of fit” was determined for the REM sleep measured from each animal (**E**) and the percentage of mice exhibiting a significant diurnal rhythm for each group plotted (**F**). The scheduled feeding increases the number of mice with rhythms in REM sleep. The amplitude of the rhythms (**G**) as well as the % of REM sleep during the night (**H**) and the total minutes of recorded REM sleep (**I**) were displayed for each group. The TRF treatment increased the % REM sleep in the night in the male BACHD but reduced the total amount of REM sleep in the female BACHD.

### NREM sleep

We examined the rhythm in NREM sleep looking at 2-hr bins across the 24-hr cycle under ALF and TRF (**Fig. 3A, B**). We analyzed the waveforms using a three-way ANOVA with genotype, treatment, and time as factors for male and female mice. For both sexes, time was highly significant and there was a significant interaction between treatment and time (**Table 1**). The reduction in NREM sleep at ZT 2 characteristic of the male BACHD was corrected by the TRF intervention. There were significant differences in the peak phase of the NREM sleep rhythms between genotypes and sexes (**Fig. 3C, D**; **Table 2**). The BACHD exhibited a great deal of variability in phase (**Table 3**) compared to the WT and this variability was reduced by TRF. For NREM sleep, this effect was even more robust in the female BACHD. The percentage of mice with a significant diurnal rhythm in REM sleep was increased by the treatment in all but the WT males (**Fig. 3E, F**; **Table 4**). The amplitude of the rhythms or the % sleep during the night did not vary with genotype, sex, or feeding schedule (**Fig. 3G, H**, **Table 2**). The total amount of NREM sleep did not vary with genotype or treatment but did exhibit sex differences (**Fig. 3I**).

**Fig 3.**
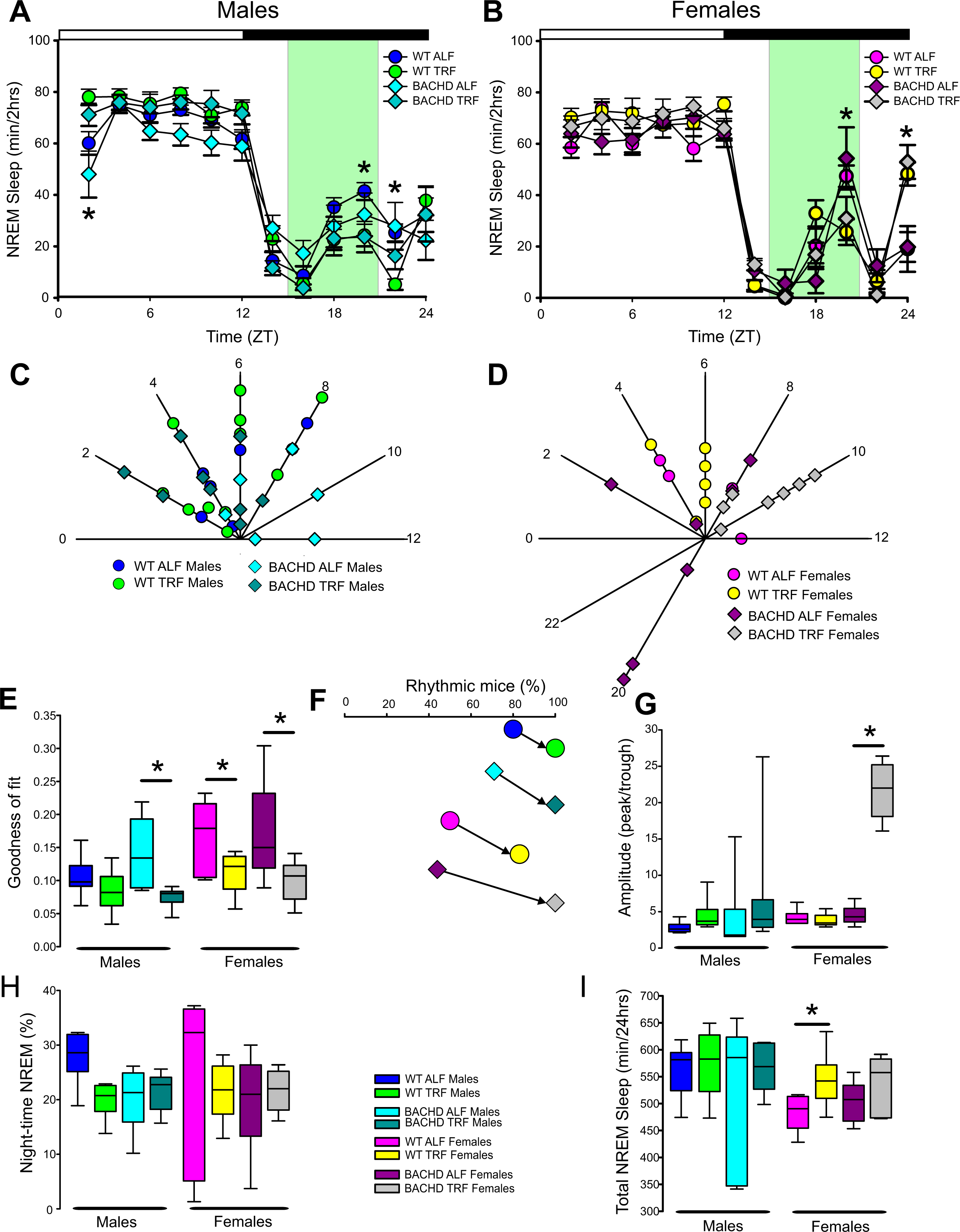
Temporal Pattern of NREM sleep in BACHD and WT mice under TRF and ALF. EEG recordings were conducted over 24-hr in undisturbed mice held in a 12:12 light/dark cycle. Waveforms showing the daily rhythms of NREM sleep in male (**A**) and female (**B**) WT and BACHD mice plotted in 2-hrs bins. The BACHD males displayed reduced NREM sleep at ZT 2 and this reduction was corrected by the TRF intervention. In WT mice, the intervention reduced NREM sleep at ZT 20. Polar display of the peak phase of the rhythms of REM sleep in male (**C**) and female (**D**) WT and BACHD mice. The female BACHD under ALF exhibited higher variability in peak phase than the other groups. The “goodness of fit” was determined for the NREM sleep measured from each animal (**E**) and the percentage of mice exhibiting a significant diurnal rhythm for each group plotted (**F**). The amplitude of the rhythms (**G**) as well as the % of NREM sleep during the night (**H**) and the total minutes of recorded NREM sleep (**I**) were displayed for each group. The TRF treatment increased the amplitude and increase the total amount of NREM sleep in the female mice.

Finally, we carried out an additional analysis by assessing sleep fragmentation by measuring the number of NREM sleep bouts and their duration using a three-way ANOVA with genotype, treatment, and sex as factors for day and night (**Fig. 4**). During the day, there were significant effects of genotype but not treatment and sex. There were also significant interactions between the three factors (**Table 5)**. Specifically, the male BACHD have more bouts than the WT and this difference is lost under TRF. During the night, there were significant effects of genotype, treatment, and sex, as well as a significant interaction between these factors (**Table 5)**. Again, the male BACHD were the most impacted and displayed a higher number of NREM sleep bouts that were of a shorter duration. TRF reduced the number of bouts to WT levels in the males.

**Fig. 4.**
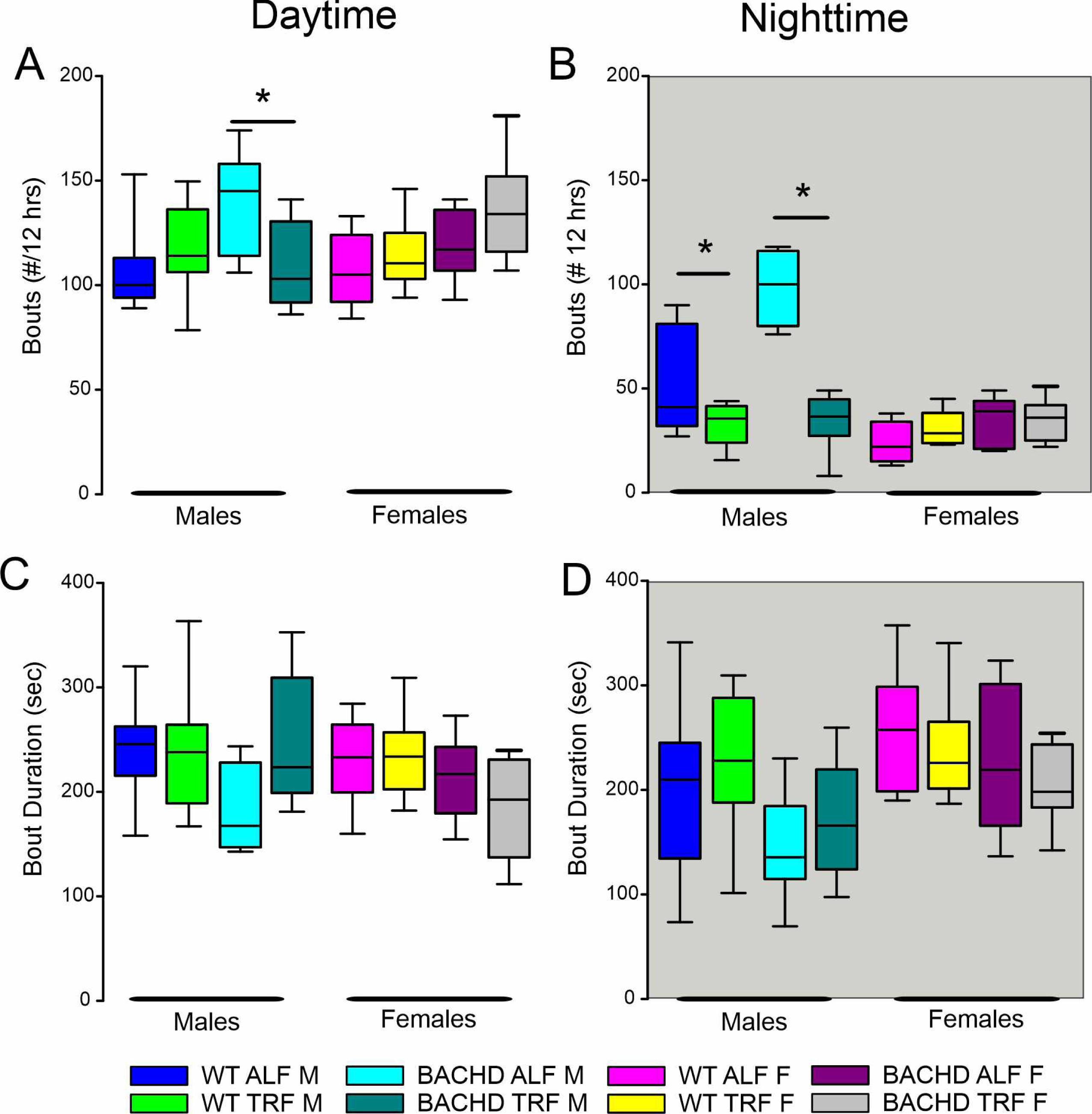
NREM sleep fragmentation in BACHD and WT mice under TRF and ALF. Sleep fragmentation during NREM sleep was calculated from the 24-hr EEG recordings. The number and average duration of the bouts were measured during the **(A, C)** day and **(B, D)** night. In the day and night, the male mutants exhibited an increased number of bouts (**A, B**) that were reduced to WT levels by the TRF intervention. No significant differences were seen with the bout duration. Significant differences between feeding schedules are indicated by an asterisk.

**Table 5:** Analysis of NREM sleep fragmentation by three-way ANOVA with genotype (WT, BACHD), sex (males, females) and treatment (ALF, TRF) as factors. Interactions between the three factors are reported. Degrees of freedom are reported within parentheses, alpha=0.05. Bold type indicates statistical significance.

| NREM |  | Genotype | Sex | Treatment | Interaction<br>(Genotype x Sex x Time) |
| --- | --- | --- | --- | --- | --- |
| Bout # | Day | **F<sub>(1, 62)</sub>= 7.306, P = 0.009** | F<sub>(1, 62)</sub>=0.032, P = 0.857 | F<sub>(1, 62)</sub>=0.062, P = 0.805 | **F<sub>(1, 62)</sub>=5.616, P = 0.021** |
|  | Night | **F<sub>(1, 62)</sub>= 15.104, P <0.001** | **F<sub>(1, 62)</sub>= 41.542, P <0.001** | **F<sub>(1, 62)</sub>= 26.766, P <0.001** | **F<sub>(1, 62)</sub>=6.361, P = 0.015** |
| Bout | Day | **F<sub>(1, 62)</sub>=5.893, P = 0.019** | F<sub>(1, 62)</sub>=1.211, P = 0.276 | F<sub>(1, 62)</sub>=0.768, P = 0.385 | **F<sub>(1, 62)</sub>=4.652, P = 0.035** |
| Duration | Night | **F<sub>(1, 62)</sub>=7.101, P = 0.010** | **F<sub>(1, 62)</sub>=9.156, P = 0.004** | F<sub>(1, 62)</sub>=0.051, P = 0.822 | F<sub>(1, 62)</sub>=0.005, P = 0.914 |

### EEG spectral power

Young BACHD mice exhibit modest changes in the EEG spectra compared to WT mice with sex differences also noted (Chiem et al., 2024). To investigate the possible impact of TRF on the EEG spectra, we quantified the power values in the frontoparietal cortical region during NREM sleep. Analysis of the power spectral curves (0.1-40Hz) with LME model found evidence for significant differences with frequency, genotype, and sex (**Table 6**). We also used a 3-way ANOVA to probe for possible impacts of the diet (**Table 7**). Treatment only altered the power spectral curves for male BACHD during the day primarily by altering the lower end of the frequency spectrum. We also examined the relative power throughout the 24-hr cycle for delta (0.5-4 Hz), theta (5-7 Hz), beta (14-20 Hz), and gamma (30-40 Hz) (**Fig. 5A-H**). The LME model found significant effects of time for delta, theta, and gamma (**Table 6**). This analysis really highlighted the broad impact of all of our factors on theta rhythms. Again, we did an additional analysis with three-way ANOVA (**Table 8**). Time exhibited a significant effect in all of the frequency bands and the females exhibited a significant interaction between time and treatment. These interactions were statistically robust and suggest a complexity in how the females responded to the treatment. We also used a linear mixed effects model to analyze this data set. Overall, apart from theta rhythms, the analysis of the spectral distribution of the EEG did not uncover robust impacts of the scheduled feeding.

**Fig. 5:**
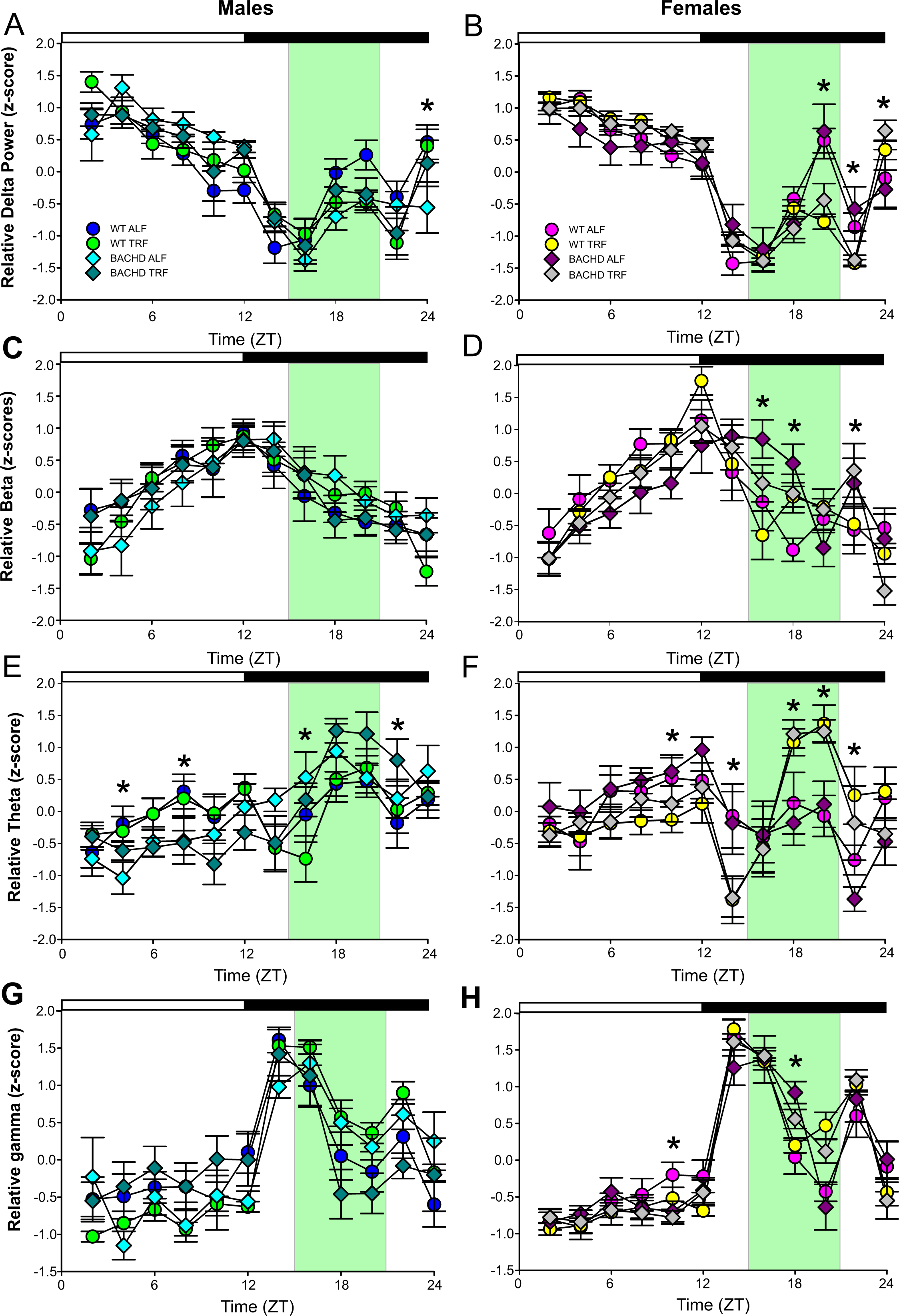
Absolute and relative power spectral analysis in BACHD and WT mice under TRF and ALF. Power spectral analysis was performed by applying a fast Fourier transform to raw 24-hr EEG waveforms. The relative power for delta (**A, B**), beta (**C, D**), theta (**E, F**) and gamma (**G, H**) power was also plotted for each group. The green shading indicates the time of feeding for the TRF groups. The symbols shown in panels A and B apply to the rest of the figure. Data are shown as the mean ± SEM. Frequencies in which treatment-evoked significant differences (P<0.05) were found with between the groups using multiple comparison procedures (Holm-Sidak method) are indicated by asterisk. The theta rhythm was most impacted by the scheduled feeding.

**Table 6:** Analysis of absolute and relative power (z-score) using a linear mixed effects (LME) model. The LME model incorporate fixed effects (genotype, treatment, sex, and time) as well as helping us understand the extent to which variability in the outcome is explained by the fixed and random effects. This analysis was not used for hypothesis testing. Bold type indicates statistical significance.

| **Absolute power** | Estimate (SE) | df | *t* | *p* |
| --- | --- | --- | --- | --- |
| Intercept | 15537.554 (80.872) | 316 | **19.012** | **< 0.001** |
| **Hz** | -64.460 (3.582) | 3116 | **-17.995** | **< 0.001** |
| **Genotype** | -302.450 (123.534) | 316 | **-2.448** | **0.014** |
| **Sex** | -570.009 (123.534) | 316 | **-4.614** | **< 0.001** |
| treatment | 25.892 (118.385) | 316 | 0.219 | 0.827 |

| **Delta** | Estimate (SE) | df | *t* | *p* |
| --- | --- | --- | --- | --- |
| **Intercept** | **0.864 (0.167)** | **764** | **5.070** | **< 0.001** |
| Genotype | 0.124 (0.252) | 764 | 0.492 | 0.623 |
| Sex | 0.328 (0.252) | 764 | 1.300 | 0.194 |
| **time** | **-0.065 (0.011)** | **764** | **-5.742** | **< 0.001** |
| treatment | 0.269 (0.252) | 764 | 1.068 | 0.286 |

| **Beta** | Estimate (SE) | df | *t* | *p* |
| --- | --- | --- | --- | --- |
| Intercept | -0.087 (0.195) | 764 | -0.445 | 0.656 |
| Genotype | 0.530 (0.296) | 764 | 1.789 | 0.074 |
| Sex | -0.159 (0.296) | 764 | -0.538 | 0.591 |
| time | 0.006 (0.013) | 764 | 0.504 | 0.615 |
| treatment | 0.158 (0.296) | 764 | 0.534 | 0.593 |

| **Theta** | Estimate (SE) | df | *t* | *p* |
| --- | --- | --- | --- | --- |
| Intercept | 0.680 (0.187) | 764 | **3.634** | **< 0.001** |
| Genotype | -0.564 (0.283) | 764 | **-1.994** | **0.046** |
| Sex | -1.654 (0.283) | 764 | **-5.845** | **< 0.001** |
| time | -0.052 (0.012) | 764 | **-4.115** | **< 0.001** |
| treatment | -0.936 (0.283) | 764 | **-3.308** | **< 0.001** |

| **Gamma** | Estimate (SE) | df | *t* | *p* |
| --- | --- | --- | --- | --- |
| Intercept | -0.895 (0.177) | 650 | **-5.068** | **< 0.001** |
| Genotype | 0.174 (0.267) | 650 | 0.672 | 0.502 |
| Sex | 0.006 (0.279) | 650 | -0.023 | 0.981 |
| time | 0.007 (0.001) | 650 | **5.739** | **< 0.001** |
| treatment | 0.002 (0.267) | 650 | -0.099 | 0.921 |

**Table 7:** Analysis of power (absolute) spectral curves during the day and the night by three-way ANOVA with genotype (WT, BACHD), sex (male, female) and frequency (1-40Hz) as factors. No significant interactions between the three factors were detected. Degrees of freedom are reported within parentheses, alpha=0.05. Bold type indicates statistical significance.

|  |  | Genotype | Frequency<br>(Hz) | Treatment |
| --- | --- | --- | --- | --- |
| Males | Day | **F<sub>(1, 809)</sub>=12.296, P<0.001** | **F<sub>(29, 809)</sub>=90.261, P<0.001** | **F<sub>(1, 809)</sub>=16.933, P<0.001** |
|  | Night | F<sub>(1, 809)</sub>=1.129, P=0.288 | **F<sub>(29, 809)</sub>=26.910, P<0.001** | F<sub>(1, 809)</sub>=0.806, P=0.370 |
| Females | Day | **F<sub>(1, 809)</sub>=28.400, P<0.001** | **F<sub>(29, 809)</sub>=96.002, P<0.001** | F<sub>(1, 809)</sub>=0.808, P=0.369 |
|  | Night | **F<sub>(1, 809)</sub>=7.498, P=0.006** | **F<sub>(29, 809)</sub>=49.516, P<0.001** | F<sub>(1, 809)</sub>=0.156, P=0.693 |

**Table 8:** Analysis of relative power (z-score) spectral curves during the day and the night by three-way ANOVA with genotype (WT, BACHD), treatment (ALF, TRF) and time as factors. Interactions between the time and treatment are reported. Degrees of freedom are reported within parentheses, alpha=0.05. Bold type indicates statistical significance.

| Frequency | Sex | Genotype | Time<br>(2 hr bins) | Treatment | Interaction<br>(treatment x Time) |
| --- | --- | --- | --- | --- | --- |
| Delta | males | F<sub>(1,431)</sub> < 0.001, P = 0.995 | **F<sub>(11,431)</sub> =35.363, P < 0.001** | F<sub>(1,431)</sub> < 0.001, P = 0.999 | F<sub>(11,31)</sub> =2.207, P = 0.089 |
|  | females | F<sub>(1,347)</sub> < 0.001, P = 0.996 | **F<sub>(11,347)</sub> =75.706, P < 0.001** | F<sub>(1,347)</sub> < 0.001, P = 0.994 | **F<sub>(11,347)</sub> =6.005, P < 0.001** |
| Beta | males | F<sub>(1,431)</sub> < 0.001, P = 0.999 | **F<sub>(11,431)</sub> =13.179, P < 0.001** | F<sub>(1,431)</sub> < 0.001, P = 1.0 | F<sub>(11,31)</sub> =0.420, P = 0.947 |
|  | females | F<sub>(1,347)</sub> < 0.001, P = 0.998 | **F<sub>(11,347)</sub> =21.467, P < 0.001** | F<sub>(1,347)</sub> < 0.001, P = 0.996 | **F<sub>(11,335)</sub> =2.755, P = 0.002** |
| Theta | males | F<sub>(1,431)</sub> < 0.001, P = 0.997 | **F<sub>(11,419)</sub> =9.471, P < 0.001** | F<sub>(1,431)</sub> < 0.001, P = 1.0 | F<sub>(11,31)</sub> =1.105, P = 0.356 |
|  | females | F<sub>(1,347)</sub> < 0.001, P = 0.999 | **F<sub>(11,347)</sub> =8.837, P < 0.001** | F<sub>(1,347)</sub> < 0.001, P = 0.998 | **F<sub>(11,335)</sub> =7.405, P < 0.001** |
| Gamma | males | F<sub>(1,431)</sub> < 0.001, P = 0.999 | **F<sub>(11,419)</sub> =30.125, P < 0.001** | F<sub>(1,431)</sub> < 0.001, P = 1.0 | F<sub>(11,31)</sub> =0.546, P = 0.871 |
|  | females | F<sub>(1,347)</sub> < 0.001, P = 0.999 | **F<sub>(11,347)</sub> =84.558, P < 0.001** | F<sub>(1,347)</sub> < 0.001, P = 0.998 | **F<sub>(11,335)</sub> =2.999, P = 0.001** |

### Recovery from sleep deprivation

The most direct test of sleep homeostatic mechanisms is to examine sleep rebound in response to SD. Therefore, we examined the 18-hr of recovery interval following 6-hrs of SD from ZT 0-6. The analysis of the waveforms for each sleep state (**Fig. 6**) using a three-way ANOVA with genotype, treatment, and time (2-hr bins) as factors (**Table 9**). The SD protocol was equally successful in all groups, all of which exhibited a sleep rebound after SD (**Fig. 6**) and time was a robustly statistically significant factor. We detected genotypic differences for wake in males and for REM sleep in females. Treatment alone did not produce any significant impact on the recovery although we did see a significant interaction between treatment and time that indicates a complexity to the sleep deprivation response. Analysis of recovery sleep between ZT 6-12, showed a strong increase in the amount of sleep in all mice. Interestingly, the groups with the TRF treatment exhibited more NREM sleep (F_(3,46)_ = 6.181, *p* = 0.001). To account for potential baseline differences in NREM sleep, we analyzed the proportion of sleep gained during recovery relative to sleep lost during SD. There were no differences between the groups (**Table 10**). The normalized relative NREM delta power was averaged across the recovery period (ZT6-24) for each animal and no difference with genotype or treatment were found (**Table 10**). There was a significant sex difference in the response. Overall, the sleep homeostatic process appeared to be functional in all groups with relatively small differences observed.

**Fig. 6.**
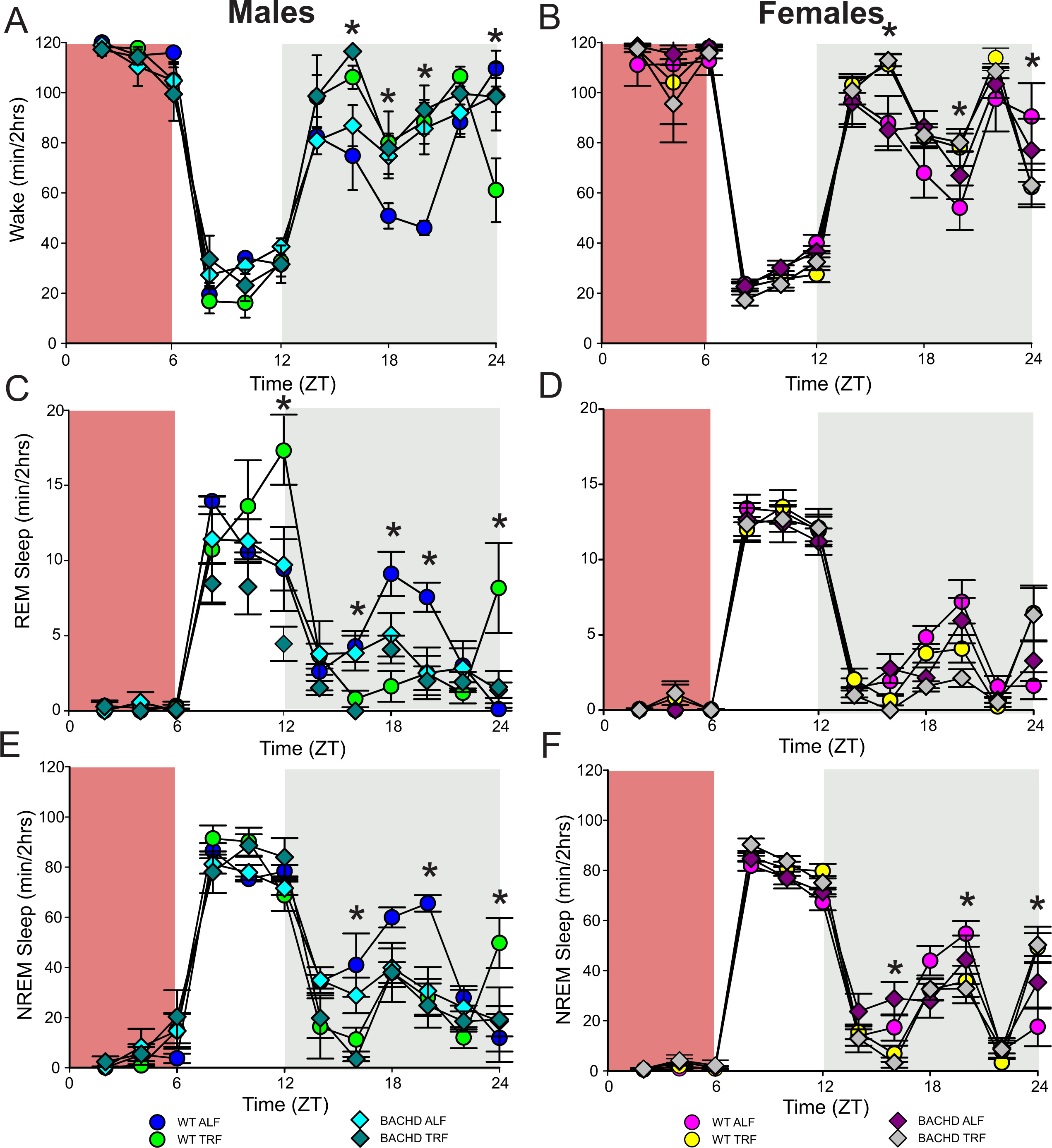
Sleep homeostatic mechanisms in BACHD mice. Mice were exposed to a 6-hr sleep deprivation (SD) at the beginning of their inactive phase (ZT0-6) using a gentle-handling protocol. EEG recordings were conducted over 24-hrs. Waveforms showing the daily rhythms in wake (**A, B**), REM sleep **(C, D)**, and NREM sleep (**E, F**) are plotted in 2-hr bins. The red-shaded area represents the time of SD. All the groups showed a strong increase in sleep between ZT6 and ZT12. Data are shown as the mean ± SEM. Time points in which significant differences (P<0.05) were found between the groups using multiple comparison procedures (Holm-Sidak method) are indicated by asterisk.

**Table 9:** Analysis of waveforms in response to sleep deprivation (SD) using three-way ANOVA with genotype (WT, BACHD), time (2 hr bins) and treatment (ALF, TRF) as factors. Interactions between the time and treatment are reported. **D**egrees of freedom are reported within parentheses, alpha=0.05. Bold type indicates statistical significance.

| State | Sex | Genotype | Time<br>(2 hr bins) | Treatment | Interaction<br>(treatment x Time) |
| --- | --- | --- | --- | --- | --- |
| Wake | males | $F_{(1,239)} = 5.406, P = 0.021$ | $F_{(11,239)} = 75.956, P < 0.001$ | $F_{(1,239)} = 4.113, P = 0.044$ | $F_{(11,239)} = 4.561, P < 0.001$ |
| | females | $F_{(1,320)} = 0.068, P = 0.795$ | $F_{(11,320)} = 120.636, P < 0.001$ | $F_{(1,320)} = 0.002, P = 0.961$ | $F_{(11,320)} = 3.840, P < 0.001$ |
| REM | males | $F_{(1,239)} = 1.141, P = 0.286$ | $F_{(11,239)} = 21.392, P < 0.001$ | $F_{(1,239)} = 2.979, P = 0.086$ | $F_{(11,239)} = 1.746, P = 0.066$ |
| | females | $F_{(1,320)} = 5.004, P = 0.026$ | $F_{(11,320)} = 160.157, P < 0.001$ | $F_{(1,320)} = 0.173, P = 0.678$ | $F_{(11,320)} = 4.415, P < 0.001$ |
| NREM | males | $F_{(1,239)} = 3.209, P = 0.075$ | $F_{(11,239)} = 71.702, P < 0.001$ | $F_{(1,239)} = 3.398, P = 0.067$ | $F_{(11,419)} = 4.209, P < 0.001$ |
| | females | $F_{(1,320)} = 0.561, P = 0.455$ | $F_{(11,320)} = 201.558, P < 0.001$ | $F_{(1,320)} = 0.005, P = 0.940$ | $F_{(11,320)} = 5.238, P < 0.001$ |

**Table 10:** Analysis of the response to SD by three-way ANOVA with genotype (WT, BACHD), sex (males, females) and treatment (ALF, TRF) as factors. No significant interactions between the three factors were detected. Degrees of freedom are reported within parentheses, alpha=0.05. Bold type indicates statistical significance.

| NREM | Genotype | Sex | Treatment |
| --- | --- | --- | --- |
| NREM sleep gained/loss | $F_{(1,48)} = 0.107, P = 0.745$ | $F_{(1,48)} = 0.261, P = 0.612$ | $F_{(1,48)} = 1.536, P = 0.222$ |
| rDelta/NREM | $F_{(1,48)} = 0.172, P = 0.680$ | $F_{(1,48)} = 14.770, P < 0.001$ | $F_{(1,48)} = 0.191, P = 0.664$ |

## Discussion

Several good models of HD have been created, each with their own advantages and disadvantages (Pouladi et al., 2013). In order to explore the intersection between circadian dysfunction and neurodegenerative disorders, we have been working primarily with the BACHD mouse model of HD. This model expresses the full-length human mutant Htt (mHtt) with 97 CAG repeats (Gray et al., 2008) and has strong construct validity as it expresses the human mutation under the control of the gene’s endogenous promoter. The face validity is also high as the BACHD model has been shown to reproduce progressive behavioral deficits such as sleep and circadian disturbances as well as selective cortical and striatal atrophies (Gray et al., 2008; Menalled et al., 2009). Of course, no animal model is perfect and BACHD mice gain body weight with age (Pouladi et al., 2012), while HD patients exhibit weight loss. Also, these mutants do not express the same transcriptional signature seen in other HD models (Gu et al., 2022).

While acknowledging these shortcomings, we feel that this mutant provides a model with strong construct and face validity. In a series of studies, we have detailed the progression of sleep, circadian, and cardiovascular deficits along with sex differences in the BACHD model (Kudo et al., 2011; Schroeder et al., 2016; Kuljis et al., 2016; 2018; Park et al., 2021; Chiem et al., 2024). These studies demonstrate that these mutants exhibit deficits in core neurodegenerative disorder-related symptoms and encourage our use of this model for circadian-based interventions.

One of the most powerful regulators of the circadian system is the daily feed/fast cycle and many studies have found benefits in scheduled feeding (Acosta-Rodríguez et al., 2022; Hepler et al., 2022; Mihaylova et al., 2023). In the present study, we used EEG to determine if scheduled feeding countered HD-driven changes in the temporal patterning of vigilance states (wake, NREM sleep, REM sleep) in the BACHD model in 6-month-old mice. We exposed the mice at the onset of their symptoms to a 6-hrs feeding/18-hrs fasting regimen aligned with the middle (ZT 15-21) of their active time (ZT 12-24). Following 1 month of treatment, we found that TRF altered several key sleep parameters with the BACHD males being particularly responsive. The scheduled feeding reduced inappropriate activity at the beginning of the day and increased the NREM sleep at the same phases. The TRF notability reduced variability in the peak phase of wake and NREM sleep in the mutants. The number of sleep bouts was notably reduced in the TRF treated mice. Our findings demonstrated clear dysfunction in the sleep/wake architecture and sleep fragmentation in adult male BACHD mice compared to WT controls and these deficits were improved by TRF treatment.

These new findings fit into a body of work where we tested whether TRF improved symptoms of HD in the BACHD (Whittaker et al., 2018) and Q175 (Wang et al., 2018) mouse models. We demonstrated that TRF-treated mutants showed improvements in locomotor activity and sleep behavioral rhythms, as well as in heart rate variability, suggesting an amelioration of the autonomic nervous system dysfunction. At the molecular level, TRF altered the phase of the rhythms of the clock gene PER2 measured both *in vivo* and *in vitro*. Importantly, TRF-treated BACHD and Q175 models exhibited improved motor performance compared with untreated mutants, and the motor improvements were correlated with improved circadian output. In the Q175 line, we found that the expression of several HD-relevant markers was restored to WT levels in the striatum of the treated mice using NanoString gene expression assays including BDNF signaling pathways.

In this earlier work, we documented TRF improvements in sleep behavior. These behavioral measures of sleep are widely used by the *Drosophila* research community (e.g. Artiushin and Sehgal, 2017) and have found acceptance by behavioral neuroscientists as well (Fisher et al., 2012; Pack et al., 2007). Using behavioral measures of sleep, we have been able to show that TRF improved the precision of sleep onset while the fragmentation of sleep was reduced. There has been a report in *Drosophila* that TRF can improve sleep behavior (Gill et al., 2015). While behavioral measures are powerful, there is a concern of using activity measures in models that are known to exhibit motor dysfunction. In rodent models, EEG determinants of sleep help address this concern about activity measures.

The possibility of sex differences in the HD phenotype has been a prominent feature of our earlier work. The male BACHD mice exhibit more severe deficits in activity rhythms and motor coordination than females (Kuljis *et al*., 2016). In recent work, we also documented sex differences in the EEG-defined sleep/wake cycle of BACHD mice at 3-months of age (Chiem et al., 2024). We found male BACHD mice exhibited reduced amounts of NREM sleep in the day, a feature which is not seen in the female BACHD. Additionally, the male BACHD mice exhibited a striking increase in variability in the phase of the rhythms in each state. While not the focus of the present study, we continued to find sex differences in vigilance states of the sleep/wake cycle of BACHD mice at 6 months of age. Overall, the EEG measures of state are consistent with our prior behavioral data indicating the male BACHD mice are more impacted than female BACHD mice. These findings raise the possibility that sex-specific factors play a role in the HD symptom progression and that hormone-based treatments may have therapeutic utility.

EEG signals are widely used to evaluate function and dysfunction in the central nervous system although the details of how the characteristic frequency bands are still an area of active research (Ibarra-Lecue et al., 2022; Andrillon and Oudiette, 2023). Neural oscillations occur in different frequency bands, each with distinct functional characteristics (Saby and Marshall, 2012). Several studies have found characteristic changes in the EEG spectra in mouse models including the R6/2 (Fisher et al., 2013; Kantor et al., 2013), the R6/1 (Lebreton et al., 2015), and Q175 (Fisher et al., 2016). When we examined absolute power, we saw clear sex differences in the lower end of the power spectrum, with BACHD females exhibiting increased power compared to the males. Statistically significant effects on genotype were seen in the males during the day and females in day and night. The TRF did produce significant changes in the males during the day but not in the other groups. Still, counter to our expectations, we did not see any impact of the diet on the individual frequency bands with the ANOVAs. The linear mixed effects (LME) model did detect highly significant effects on the theta band. Given the association of hippocampal theta rhythms with memory formation and navigation, these TRF driven changes should be explored in future studies.

To test sleep homeostatic mechanisms, we examined how the BACHD and WT mice responded to a 6-hr SD protocol. All groups were effectively sleep deprived and all groups responded to SD with an increase in sleep amount. There were no differences in the amount of sleep gained during recovery relative to sleep lost during SD or in the normalized relative NREM delta power during the recovery period. Scheduled feeding alone did not produce any significant impact on the recovery. We did see a significant interaction between treatment and time that indicates a complexity to the sleep deprivation response that would require additional experiments to clarify.

The mechanisms underlying these benefits of TRF on central nervous system function are not known although there are several possibilities (Longo and Panda, 2016; Jensen et al., 2020; Bass, 2024). In recent work, we reported that a ketogenic diet (KD) ameliorates the symptoms and delays disease progression in the BACHD model (Whittaker et al., 2022). Mutant mice fed a KD for three months displayed increased daytime sleep and improved timing of sleep onset. In addition, KD improved activity rhythms and motor performance on the rotarod and challenging beam tests. Prior work has also shown benefits of the KD in the R6/2 model in reducing weight loss and improving open field behavior (Chen et al., 2016). It is noteworthy that the probiotic *Akkermansia municiphila* (*Akk*) dramatically increased in abundance under KD, since it has been associated with improved metabolic health and lower inflammation (Attaye et al., 2021; Bonnechere et al., 2022). Intriguingly, work in a *Drosophila* model of HD implicated the gut bacteria in the regulation of HD pathology (Chongtham et al., 2022), and diet-driven changes in the microbiome could also influence HD-pathology in the fly. In a project with Dr. Desplats’s group, we recently exposed Alzheimer’s disease mouse models to the TRF protocol and reported that this regimen had the remarkable capability of simultaneously reducing amyloid deposition, increasing Aβ42 clearance, and normalizing daily transcription patterns of multiple genes (Whittaker et al., 2023). Together, these findings suggest that feeding schedules and diet could play an important role in the development of new treatment options for HD and other neurodegenerative disorders.

